# Cyclophosphamide immunosuppressed Xid mice model clarify the protective role of B cells in experimental encephalitozoonosis

**DOI:** 10.1101/430256

**Authors:** Carla Renata Serantoni Moysés, Lidiana Flora Vidôto da Costa, Elizabeth Cristina Perez, José Guilherme Xavier, Diva Denelle Spadacci-Morena, Paulo Ricardo Dell’Armelina Rocha, Anuska Marcelino Alvares-Saraiva, Maria Anete Lallo

## Abstract

*Encephalitozoon cuniculi* is an intracellular pathogen that stablishes a balanced relationship with immunocompetent individuals, which is dependent of T lymphocytes activity. We previously showed X-linked immunodeficiency (XID – B cell deficient) mice are more susceptible to encephalitozoonosis and B-1 cells presence influences in the immune response. Because XID mice are deficient both in B-1 and B-2 cells, here we investigate the role of these cells against *E. cuniculi* infection using cyclophosphamide (Cy) immunosuppressed murine model to exacerbate the infection. XID mice presented lethargy and severe symptoms, associated with encephalitozoonosis and there was an increase in the peritoneal populations of CD8^+^ and CD4^+^ T lymphocytes and macrophages and also in the proinflammatory cytokines IFN-γ, TNF-α and IL-6. In BALB/c mice, no clinical signs were observed and there was an increase of T lymphocytes and macrophages in the spleen, showing an effective immune response. B-2 cells transfer to XID mice resulted in reduction of symptoms and lesion area with increase of B-2 and CD4^+^ T populations in the spleen. B-1 cells transfer increased the peritoneal populations of B-2 cells and macrophages and also reduced the symptoms. Therefore, the immunodeficiency of B cells associated to Cy immunosuppression condition leads to disseminated and severe encephalitozoonosis in XID mice with absence of splenic immune response and ineffective local immune response, evidencing the B-1 and B-2 cells role against microsporidiosis.

**Author summary:** The adaptive immune response plays a key role against *Encephalitozoon cuniculi*, an opportunistic fungus for T cells immunodeficient patients. The role of B cells and antibody play in natural resistance to *Encephalitozoon cuniculi* remains unresolved. Previously, we demonstrated that B-1 deficient mice (XID), an important component of innate immunity, were more susceptible to encephalitozoonosis, despite the increase in the number of CD4^+^ and CD8^+^ T lymphocytes. In order to better understand the role of B-1 and B-2 cells and the relationship with the other cells of the immune response in encephalitozoonosis, we infected with *E. cuniculi* in cyclophosphamide immunosuppressed mice. Here we demonstrate that infected XID mice showed reduction of T cells and macrophages and increase of proinflammatory cytokines associated with disseminated and severe encephalitozoonosis with presence of abdominal effusion and lesions in multiple organs. This pattern of infection observed in mice with genetic deficiency in T cells, so we suggest that the absence of B-1 cells affects the cytotoxic capacity of these lymphocytes. When we transfer B-2 cells to XID mice, the lesion areas caused by the fungus, the populations of T lymphocytes in the peritoneum and the proinflammatory cytokines decrease, indicating a better resolution of the infection. We speculate that B-1 and B-2 cells participate in the immune response against *E. cuniculi*, interacting with the other components effective in immunity. The results shown here indicate that B-1 cells as a constituent of the innate response to microsporidia.

## Introduction

Microsporidia are obligate intracellular pathogens belonging to Microsporidia Filum and Fungi Kingdom. These eucariotic microrganisms are capable of infecting vertebrates and invertebrates [1]. From more than 1.200 Microsporidia species so far described, the pathogens belonging to the Genus *Encephalitozoon* have been associated with mammal and human disease, especially *E cuniculi, E. intestinalis* and *E. hellem*. Ultrastructural observations showed that due to an injection structure – the polar tubule – Microsporidia are capable of infecting different types of cells, by injecting its infectious content – the sploroplasm – into the cytoplasm of infected cells [2,3].

The most affected human populations include all those immunosuppressed, mostly individuals with lymphocytic populations decrease. These populations include mostly HIV infected and patients in chemotherapy [4,5]. In decreased immune response, the pathogen development and spreads are facilitate determining cells death [6,7,8,9]. However, a previous study in mice infected with *E. cuniculi* showed that even in animals treated with albendazole the pathogen could reinfect, characterizing a latent infection [10]. Then, there is no effective treatment to *E. cuniculi*.

Cellular immunity is critical for survival of *E. cuniculi* infected host, and CD8^+^ T lymphocytes are crucial for the targeted death of infected cells, avoiding pathogen dissemination [11,12,13,14]. Depending the infection route, CD4^+^ T lymphocytes are required for efficient immune response against *E. cuniculi*, as showed in oral infection [9]. Although B cells produce antibodies against *E. cuniculi*, its effective role may be incipient and not enough for avoiding the pathogen from progressively infecting other cells [15,16]. Therefore, the role of B cells needs better investigation.

B lymphocytes are classified in B-1 cells, localized mostly in the peritoneal and pleural cavities, and B-2 lymphocytes, localized mostly in the follicle of lymph nodes, marginal zone of the spleen and bone marrow [17,18,19]. Also, these two cell subtypes have different surface cell markers, antibodies repertoire, morphology and function [20,21]. XID mice have a damage of Bruton’s Tirosine cinase (Btk) which allows B-1 and B-2 cells deficiency. These mice have been used as a model to better investigate the roles of B cells in immunity [22,23]. A previous study from our group showed that XID mice are more susceptible to encephalitozoonosis than their WT background BALB/c mice. The susceptibility was associated with B-1 cells deficiency, since the adoptive transfer of B-1 cells to XID mice increased the numbers of immune system cells and more resistance to infection by oral and peritoneal route [8,9].

We previously demonstrated that Cy causes immunosuppression and became BALB/c mice more susceptible to disseminated and lethal encephalitozoonosis [24,25]. Herein, we used Cy treatment to mimic immunocompromised conditions to better understand the immune response. After Cy immunosuppression, BALB/c and XID mice were infected with *Encephalitozoon cuniculi*. Instead the Cy immunosuppressive effects, *E. cuniculi* infection determined an increase of peritoneal macrophages and T lymphocytes and proinflammatory serum cytokines in infected XID mice. In infected BALB/c mice we observed increase of CD4^+^ and CD8^+^ T lymphocytes and macrophages into the spleen, evidencing recruitment of immune response which resulted in attenuated infection. These results help us to clarify the different roles of B-1 and B-2 cells against encephalitozoonosis.

## Results

### XID mice are more susceptible to *E. cuniculi* infection

Clinically, difficulty for movement ruffed coat, as well as serous-bloody effusion in the abdomen, hepatomegaly, splenomegaly and extended intestinal segments were observed in infected XID mice (Fig 1A). XID mice adopted transferred with B-1 and B-2 cells with few clinical signs e discrete peritoneal effusion (Fig 1B). No symptoms were observed in BALB/c mice. Peritoneal lavage counting of spores was similar between all mice groups (Fig.1B). Microscopically, lesions were observed in the liver, lungs, spleen and small intestine. In the liver we observed granulomatous lesions with multifocal mononuclear inflammatory infiltrate. At 14 DPI, in BALB/c the lesions areas are increased then XID mice (Fig. 1C). The lesion area decreases significantly after B-2 cells adoptive transfer (XID+B-2) while increase after B-1 cells adoptive transfer (XID+B-1) (Fig. 1C). Although the lesions area was decreased in XID mice, the liver presented degenerative lesions with the presence of vacuoles, picnotic nuclei and spores. Inflammatory infiltrates were observed mostly at the periportal space, as well as in the liver tissue adjacent to the subcapsular space associated with spores (data not shown). *E. cuniculi* spores were observed in the liver associated with the cells infiltrate (Fig. 2A, B, C e D). There was keratin expression in hepatocytes and others epithelial hepatic parenchyma, however, in the inflammatory focus there was no keratin reactivity, despite the granulomatous aspect of the lesions (Fig. 2E e F).

**Figure 1.**
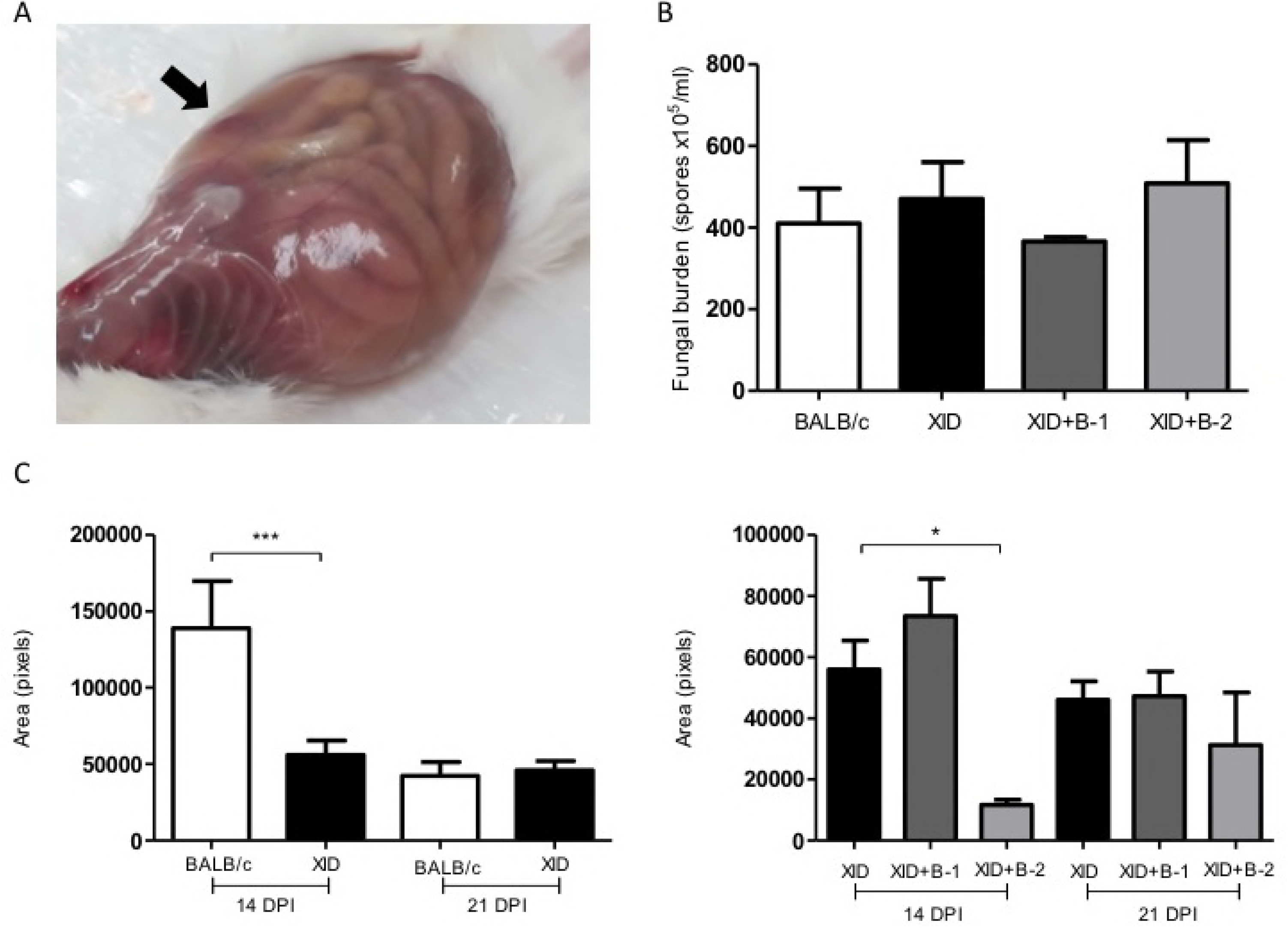
Fungal burden. A) Serous-bloody effusion in the abdominal cavity in XID mice infected with *E. cuniculi*. B) Evaluation of the spores from peritoneal washes in BALB/c, XID, XID+B-1 (adoptively transferred with B-1 cells) and XID+B-2 (adoptively transferred with B-2 cells) mice infected with *E. cuniculi*. One-way ANOVA analysis with Bonferroni multiple comparisons post-test did not reveal significance. C) Comparison of the means of the lesion areas in the livers of BALB/c, XID, XID+B-1 e XID+B-2 mice, in pixels, measured by morphometric analysis. One-way ANOVA analysis with Bonferroni multiple comparisons post-test revealed *p<0.5; *** p<0.001.

**Figure 2.**
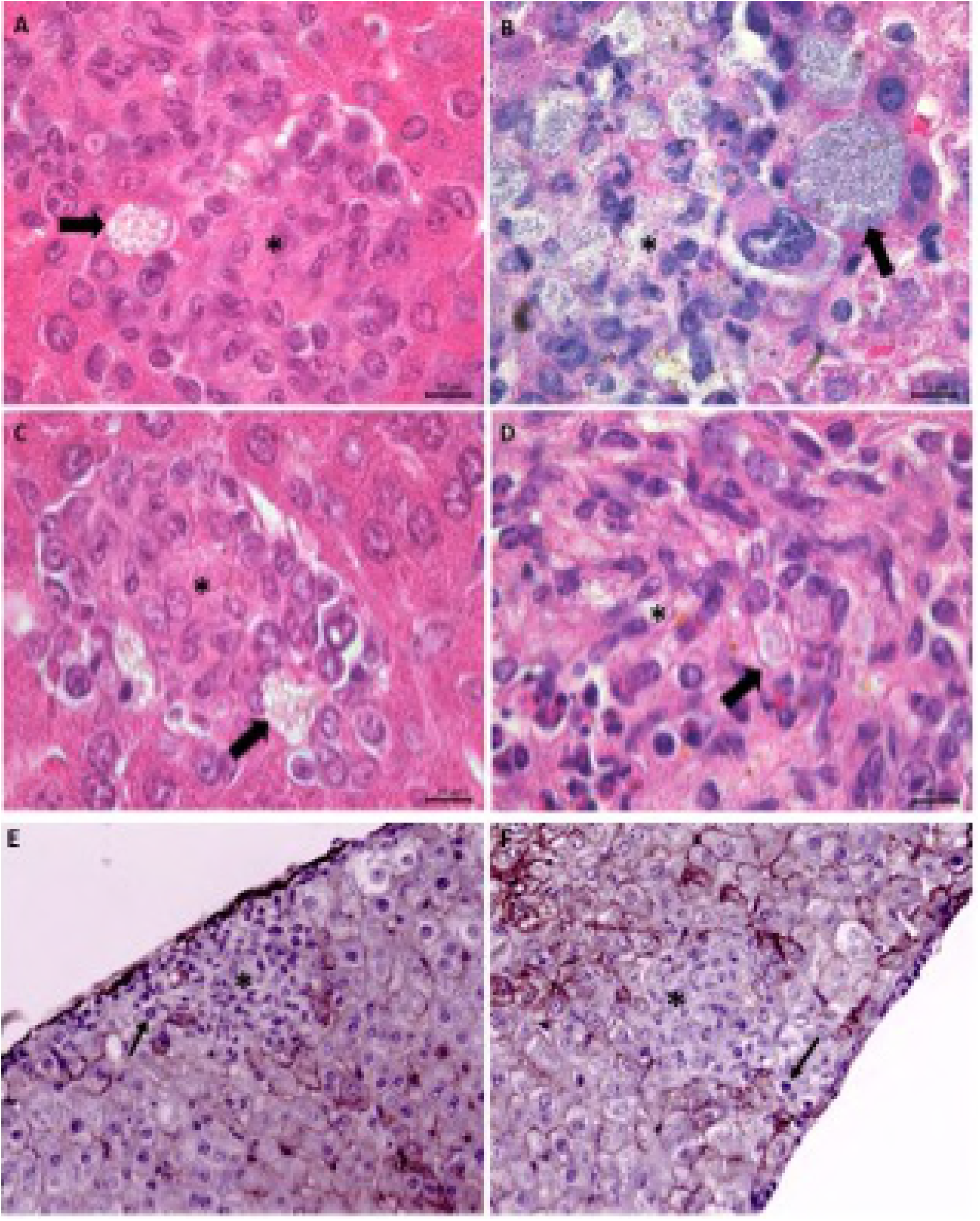
Photomicrography of hepatic lesions of mice infected with *E*. cuniculi. Spores (arrow) associated with inflammatory infiltrates in BALB/c (A), XID (B), XID+B-1 (C) e XID+B-2 (D) mice, HE stain. (E and F) Expression of keratins in the hepatic parenchyma in XID mice, but no marking in inflammatory infiltrate cells (*) and the presence of macrophages (thin arrow) associated or not with mitosis.

Transmission electron microscopy (TEM) confirmed the predominance of macrophages in the mononuclear infiltrates from XID liver with fungal spores in lysis and parasitophorous vacuoles (Fig. 3A and B), where different stages of parasite development were observed, such as meronts, sporoblasts and mature spores (Fig. 3C). Intact spores were also observed in the cytoplasm of hepatocytes and in the intercellular spaces, with evidence of the polar tubule (Fig. 3D). This result evidence the development of the pathogen into the tissue spite of the presence of immune cells. Also, we observed collagen near the lesion areas (Fig. 3B).

**Figure 3.**
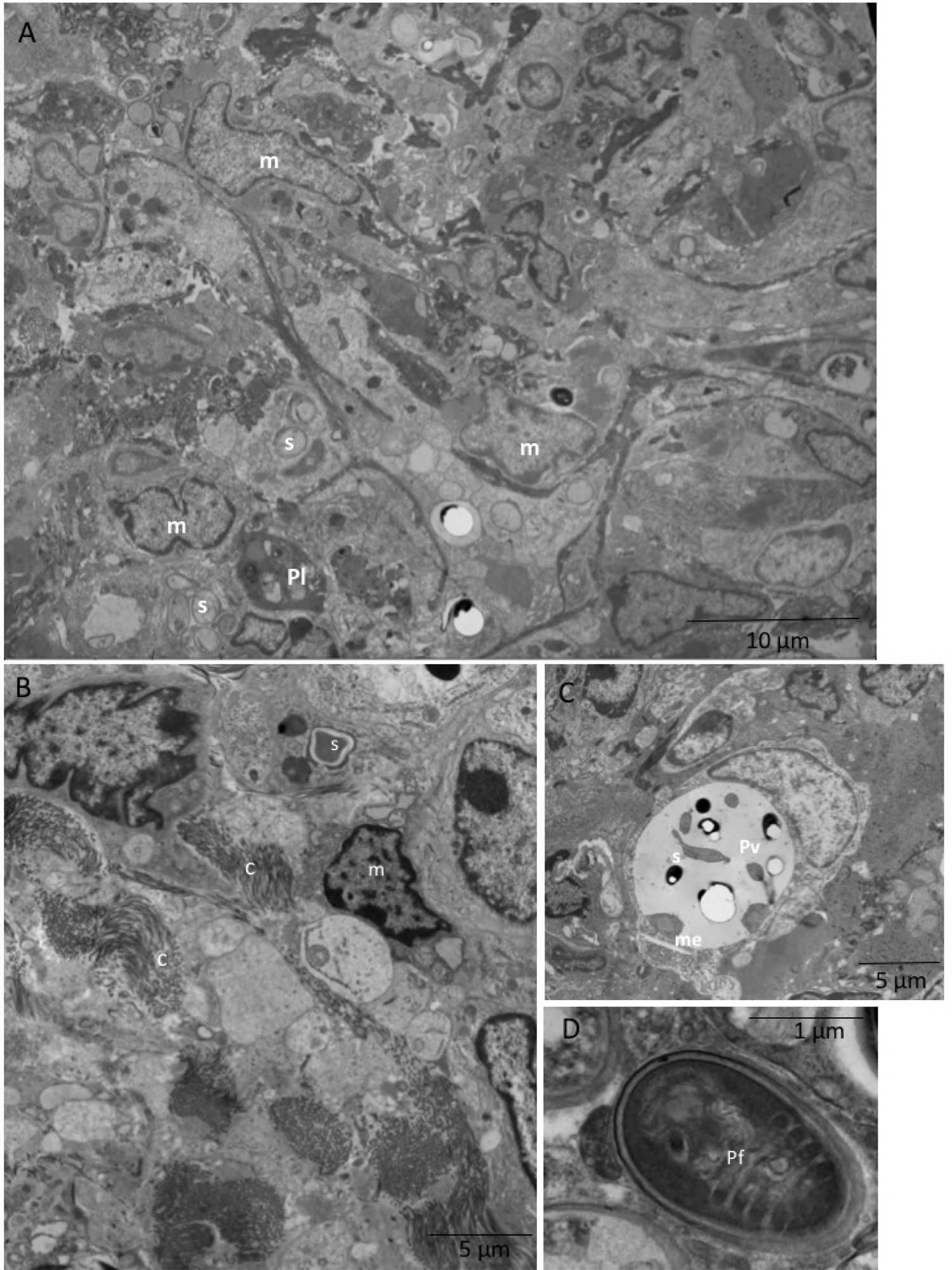
Electron micrograph of liver from XID mice treated with cyclophosphamide and infected with *E. cuniculi*. (A) Inflammatory infiltrate with predominant macrophages (m) with phagolysosome vacuoles (Pl) containing remains of spores or intact spores (e). (B) Inflammatory infiltrate with deposition of collagen (c), spores (e) and macrophages (m). (C) Parasitophorous vacuoles (Pv) with proliferative forms such as meronts and (me) e spore (s). (D) detail of a free spore in the tissue parenchyma, note the polar filament (Pf).

Multifocal interstitial pneumonia was observed in infected mice, associated with mononuclear predominantly lymphoplasmacytic infiltrate and alveolar collapse (Fig. 4). Also, lymphoplasmacytic mild enteritis was also observed in all infected groups and spores were rare (Fig. 5B).

**Figure 4.**
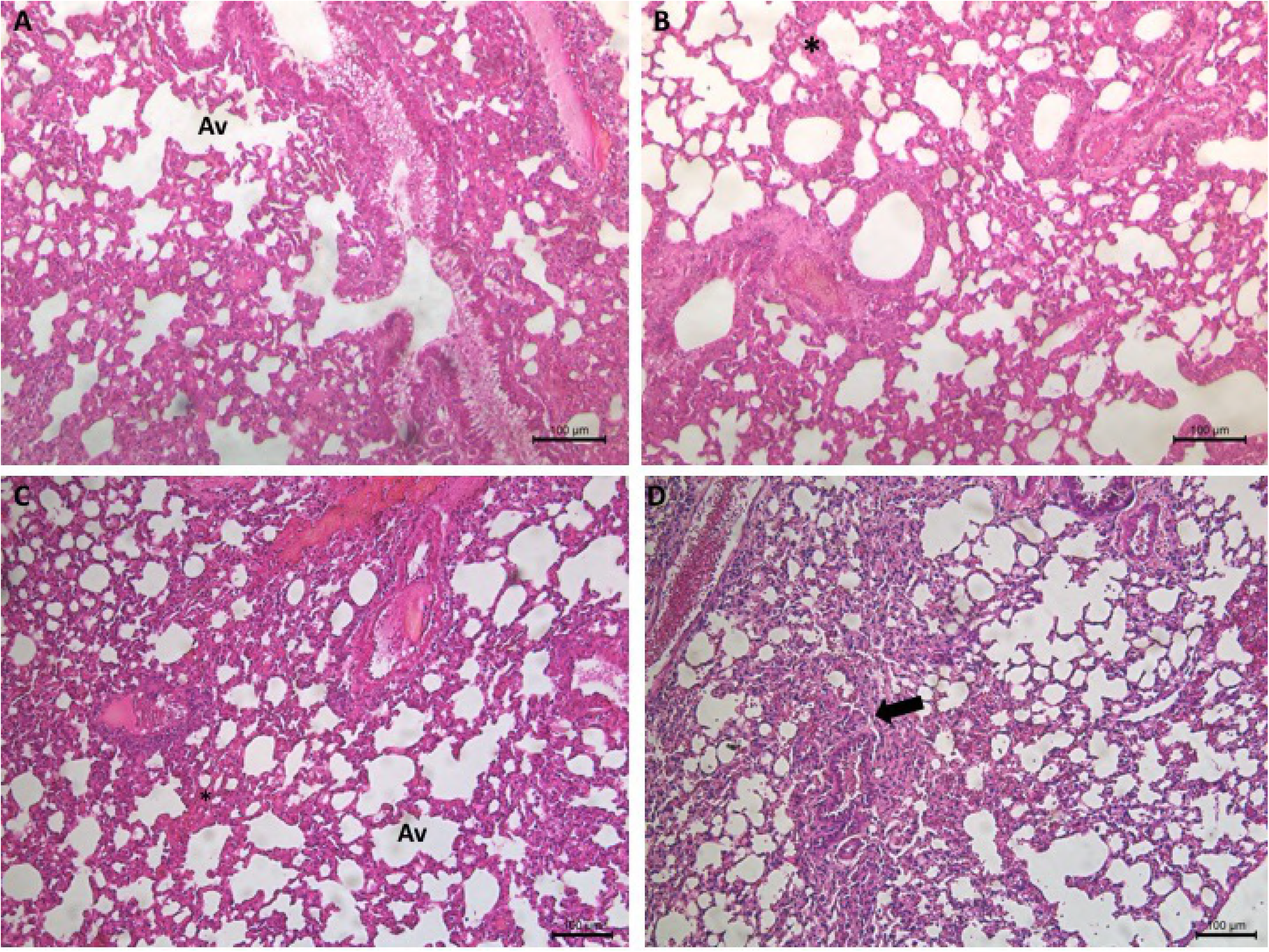
Interstitial pneumonia in infected mice. Photomicrography evidenced lymphoplasmacytic leukocyte infiltration determining walls thickening around alveoli (Av), thickened interalveolar septa (*) and areas of alveolar collapse (arrow) in BALB/c (A), XID (B), XID+B-1 (C) e XID+B-2 (D) mice. HE stain.

**Figure 5.**
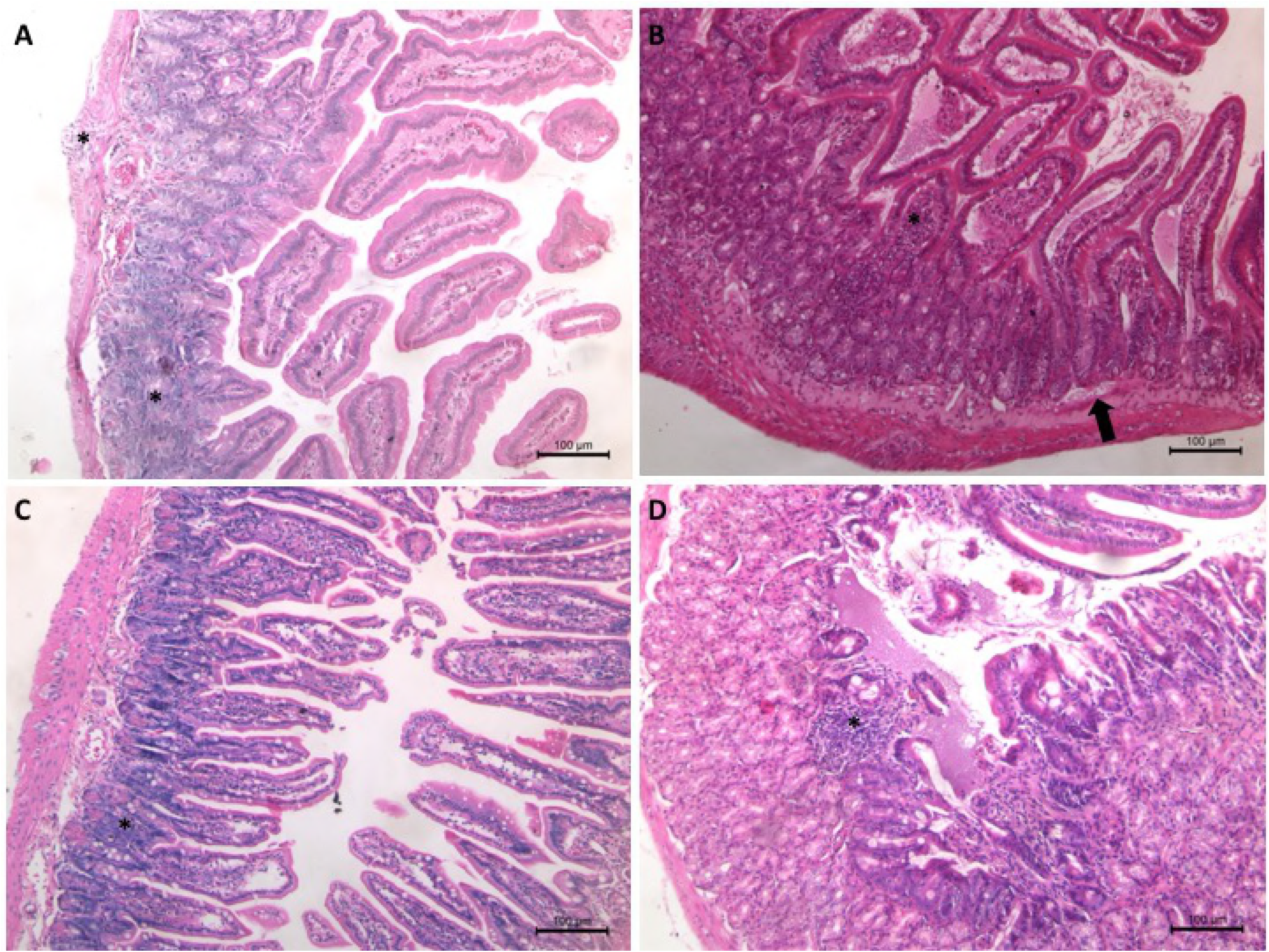
Lymphoplasmacytic enteritis in infected mice. Photomicrography of the small intestine with preserved villi and inflammatory infiltrate (*) in lamina propria, concentrated around the crypts in BALB/c (A), XID (B), XID+B-1 (C) e XID+B-2 (D) mice. *E. cuniculi* spores (arrow) in XID mice (B). HE stain.

In the spleen of infected animals, rarefaction of the white pulp was observed in BALB/c, XID and XID+B-1 (Fig. 6A, B e C) mice. Interestingly, in the spleen of only XID+B2 mice, there was an important lymphocytic infiltrate (Fig. 6D). TEM showed parasitophorous vacuoles in splenic cells from XID mice, where different development stages of *E. cuniculi* spores were observed, such as meronts in binary division, sporoblasts and sporonts (Fig. 7), reinforcing the development of the pathogen even in lymphoid tissue.

**Figure 6.**
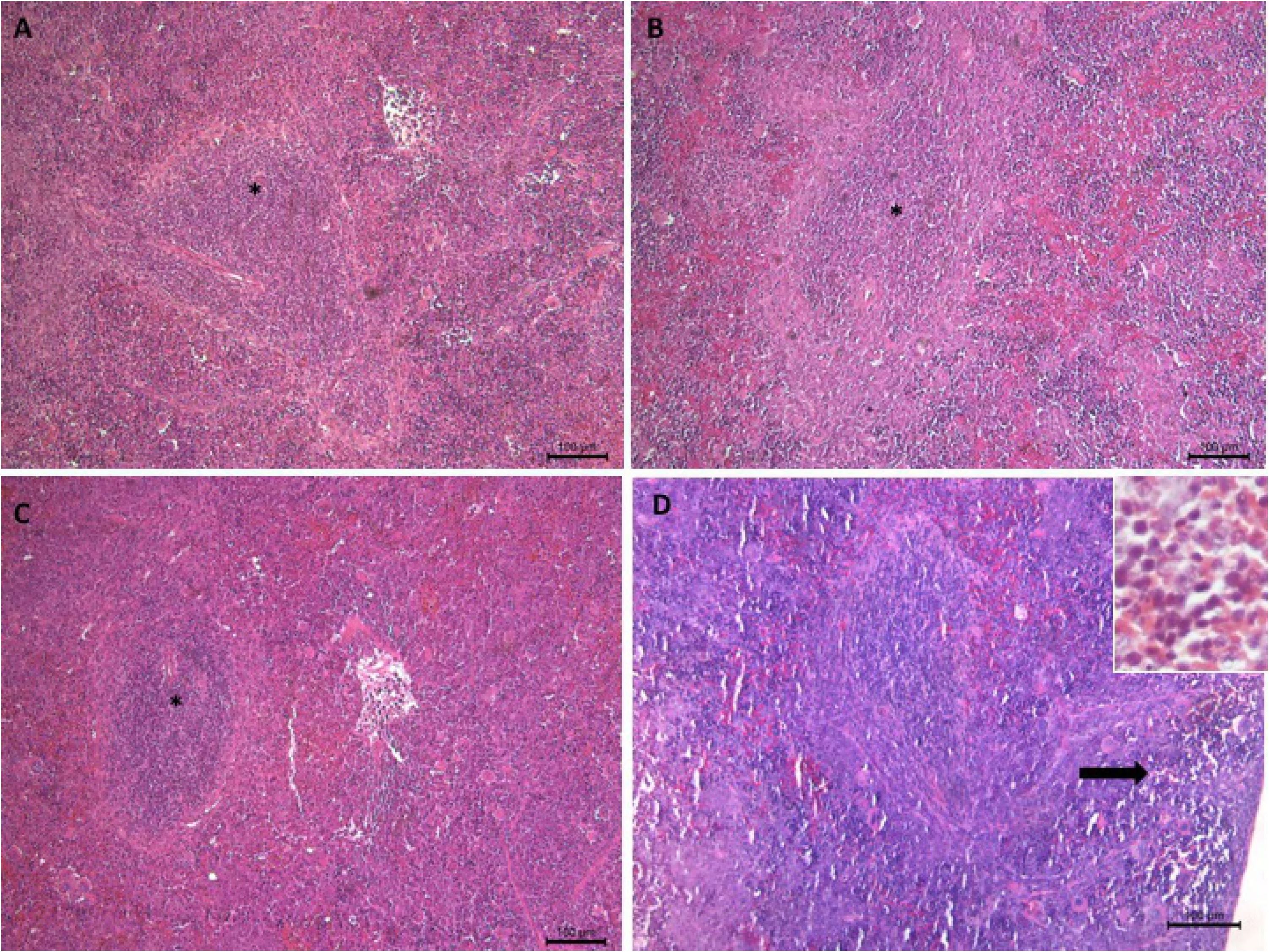
Spleen photomicrography of infected mice. Rarefaction in lymphoid white pulp (*) in BALB/c (A), XID (B) and XID + B-1 (C). Lymphocyte infiltration (arrow) in red pulp region in XID+B-2 (see detail in insert) (D). HE stain

**Figure 7.**
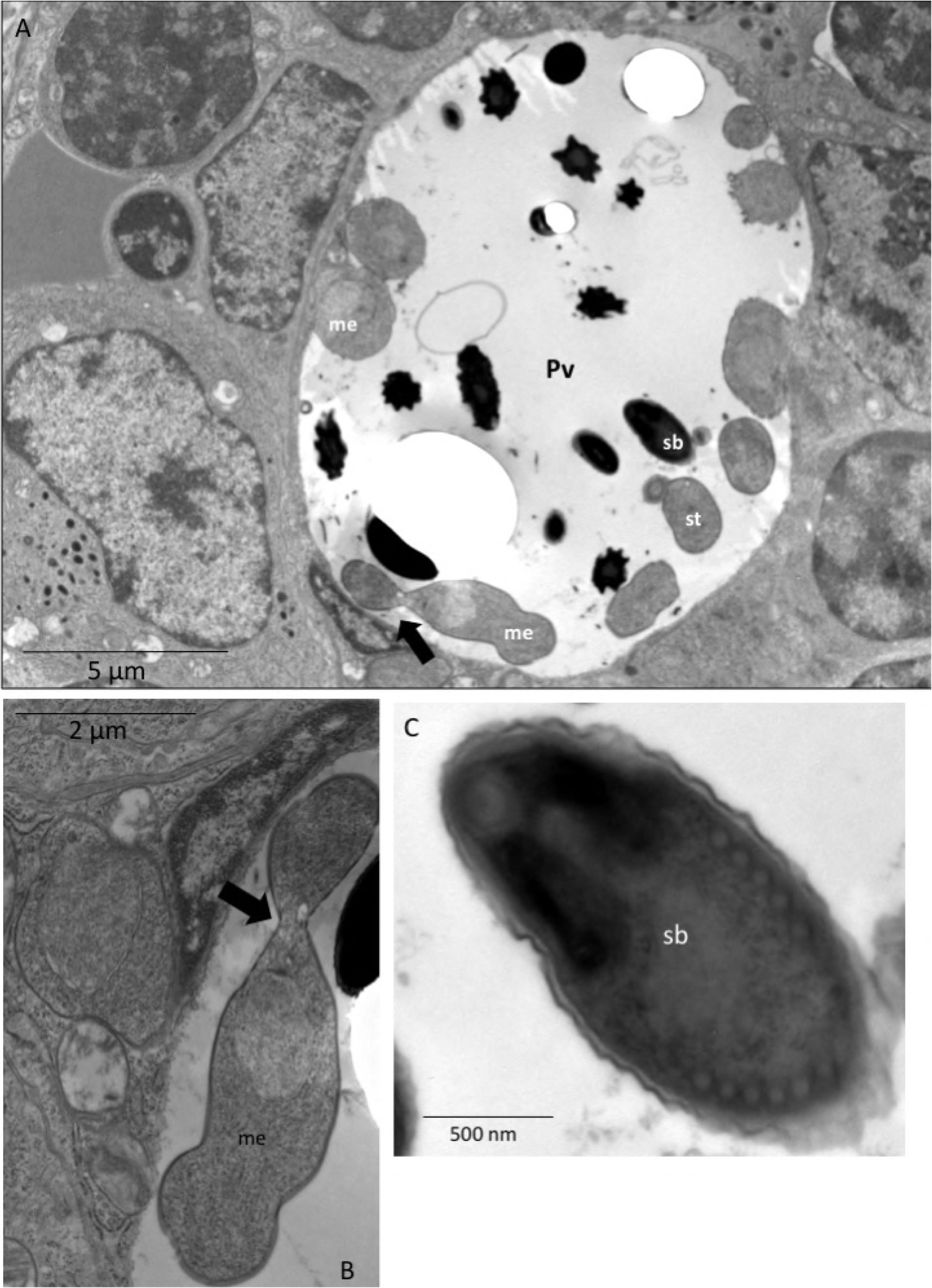
Electron micrograph of parasitophorous vacuoles with proliferative forms in the spleen of infected XID mice. (A) Parasitophorous vacuoles (Pv) with meronts (me) in the periphery, in binary fission (arrow) or not, sporoblasts (sb) and spores (s). (B) Detail of binary fission of meronte. (C) Detail of sporoblast.

### Increase of immune cells in the peritoneal cavity of infected XID mice

Phenotypical analysis of immune cells showed that CD8^+^ and CD4^+^ T lymphocytes and macrophages increased in infected XID mice compared to uninfected control (Fig. 8). In infected BALB/c mice, at 14 DPI there was an increase in the macrophage population in the peritoneum compared to uninfected. However, this population decreased at 21 DPI (Fig. 8). B-2 cells population also decreased in infected BALB/c mice. Overall, both at 14 and 21 dpi, the populations of peritoneal CD8^+^ and CD4^+^ T and B-2 lymphocytes and macrophages increased in infected XID compared to infected BALB/c mice (Fig. 8).

**Figure 8.**
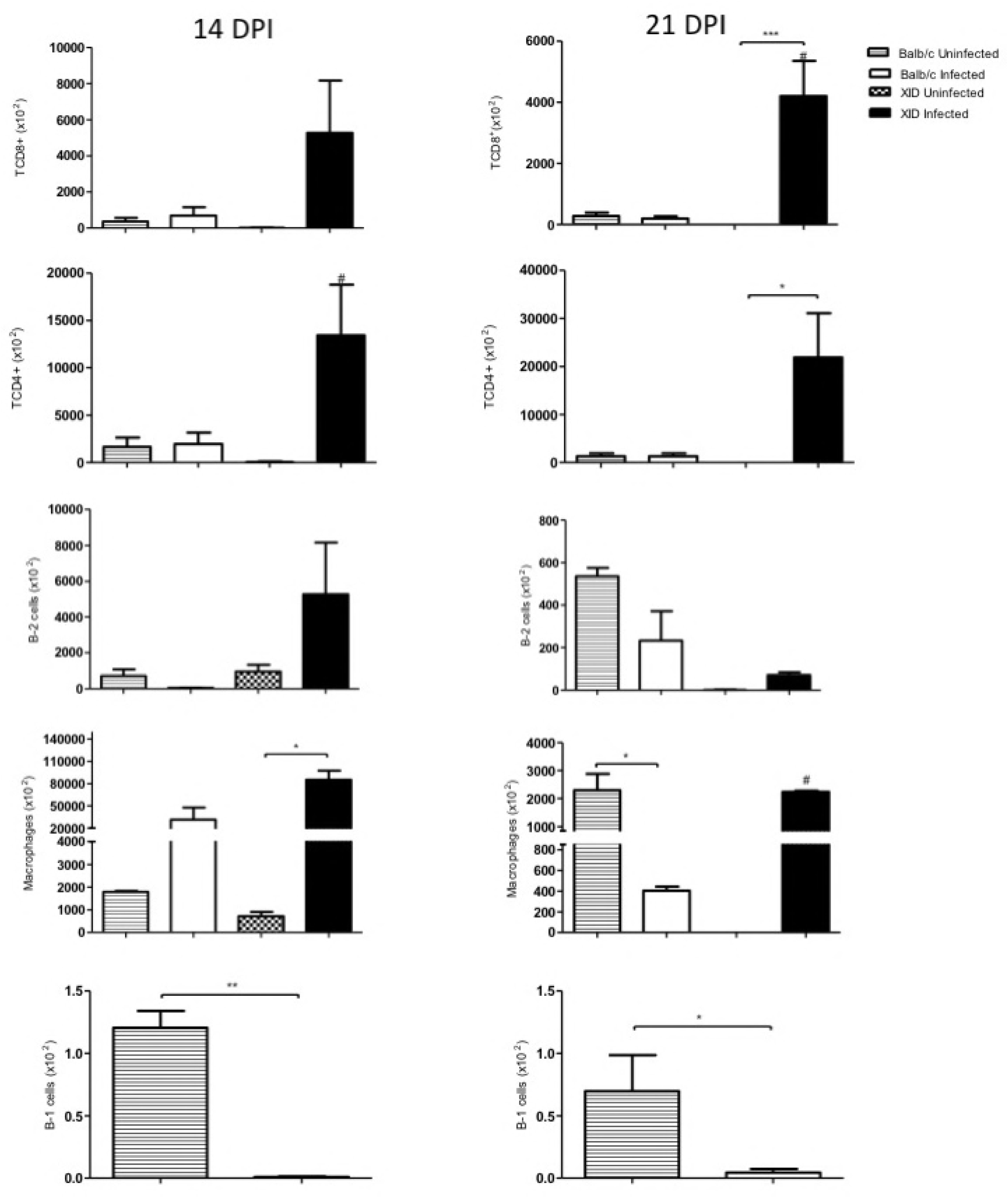
Evaluation of the peritoneal populations of CD8+ and CD4+ T lymphocytes, B-2 cells and macrophages of BALB/c and XID mice immunosuppressed with cyclophosphamide and infected or not with *E. cuniculi* at 14 or 21 days post-infection (DPI). One-way variance analysis ANOVA with Bonferroni multiple comparisons post-test showed *p<0.5; ***p<0.001 and t test unpaired #p<0.5 compared to infected BALB/c. For B-1 cells, unpaired t-test analysis revealed *p <0.5; **p <0.01.

Moreover, there was a decrease of CD8^+^ and CD4^+^ T lymphocytes in XID+B-2 mice group compared to other XID group (XID and XID+B-1) (Fig. 9). However, adoptive transfer of B-1 cells to XID mice (XID+B-1 group) caused a significant increase of B-2 cells and macrophages compared to other XID mice (Fig.9).

**Figure 9.**
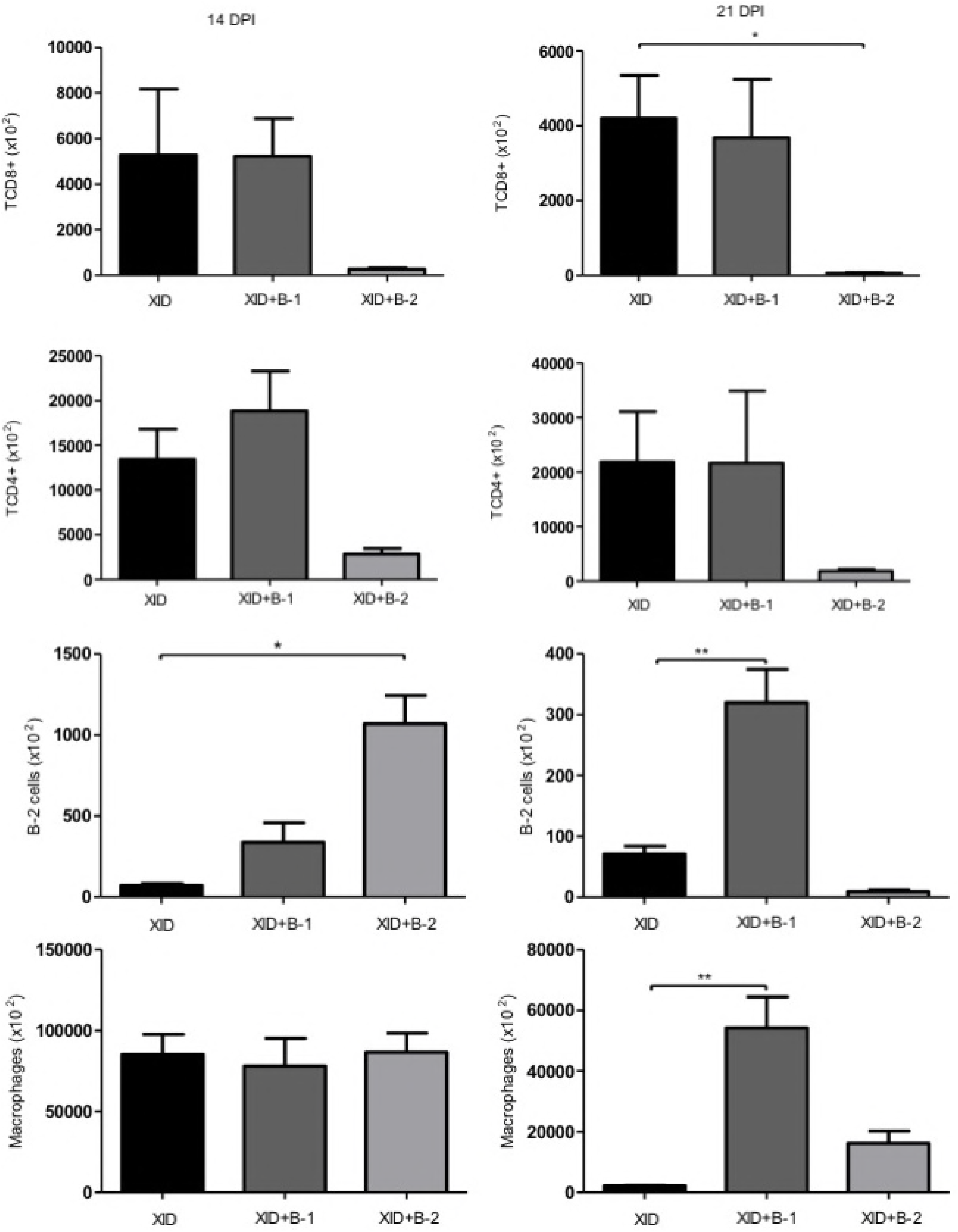
CD8^+^ and CD4^+^ T lymphocytes, B-1 and B-2 cells and macrophages in the peritoneal cavity from XID, XID+B-1 and XID+B-2 mice immunosuppressed with cyclophosphamide and infected with *E. cuniculi* at 14 and 21 days post-infection (DPI). One-way variance analysis ANOVA with Bonferroni multiple comparisons post-test, *p <0.5; **p <0.01.

### Increased immune response into spleen of BALB/c mice

In the spleen of infected BALB/c mice, there was an increase of CD8^+^ and CD4^+^ T lymphocytes and macrophages compared to uninfected control (Fig. 10). In XID mice, there was no difference in the populations of immune cells compared to uninfected control, except for B-2 cells, that decreased in XID mice at 14 DPI (Fig. 10). These results indicate important differences in the immune response assembly from spleen of XID and BALB/c mice, which could be related to the development of *E. cuniculi* in XID mice. Moreover, there was a decreased percentage of macrophages, and an increase in CD4^+^ T and B-2 lymphocytes in the spleen of XID+B-2 mice compared to XID and XID+B-1 groups (Fig. 11).

**Figure 10.**
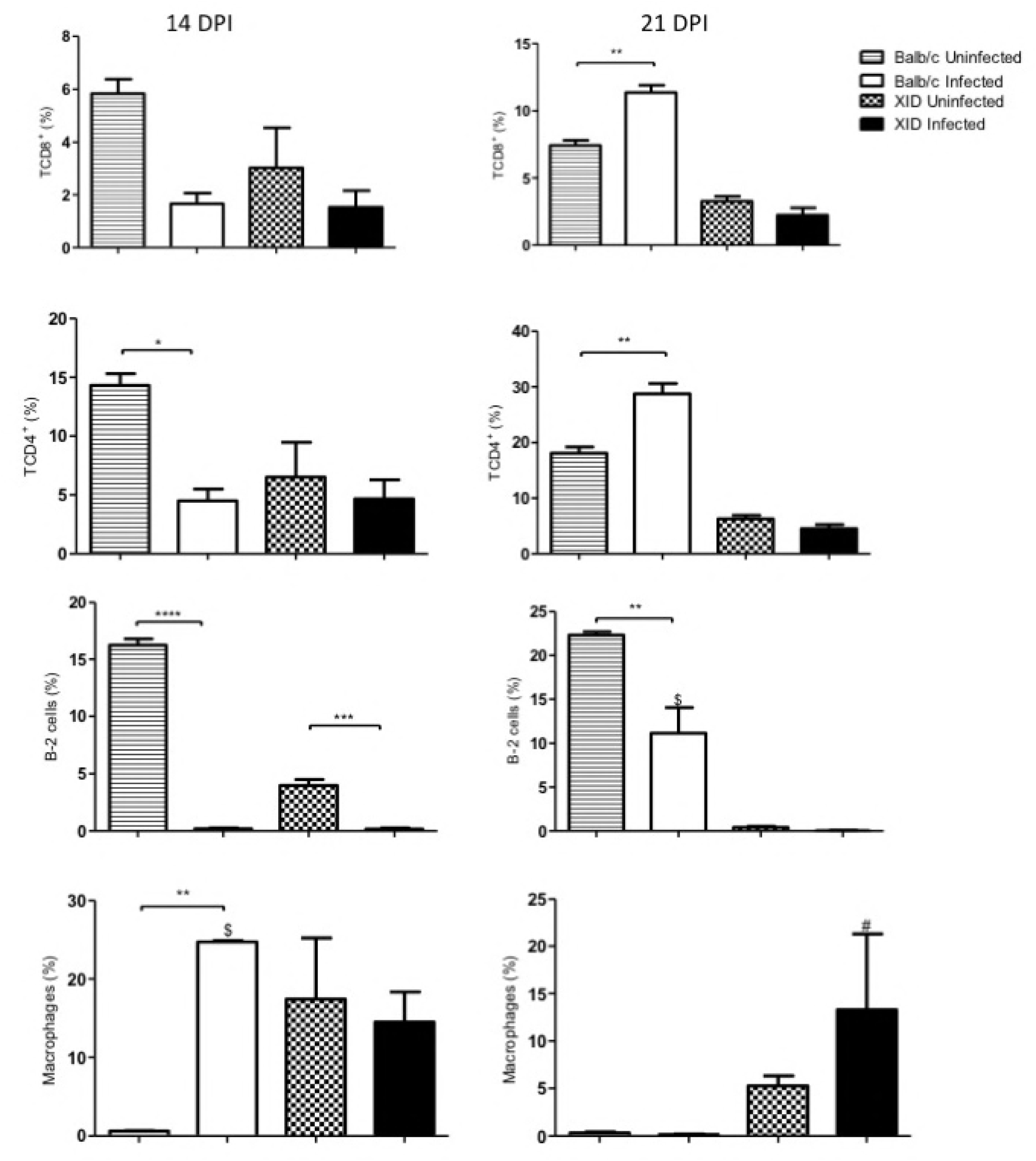
Evaluation of the splenic populations of CD8^+^ and CD4^+^ T lymphocytes, B-2 cells and macrophages of BALB/c and XID mice immunosuppressed with cyclophosphamide and infected or not with *E. cuniculi* at 14 and 21 days post-infection (DPI). One-way variance analysis ANOVA with Bonferroni multiple comparisons post-test, *p <0.5; **p <0.01; ****p <0.0001. Unpaired t-test, $p <0.01 compared to infected XID, #p <0.0001 compared to BALB/c.

**Figure 11.**
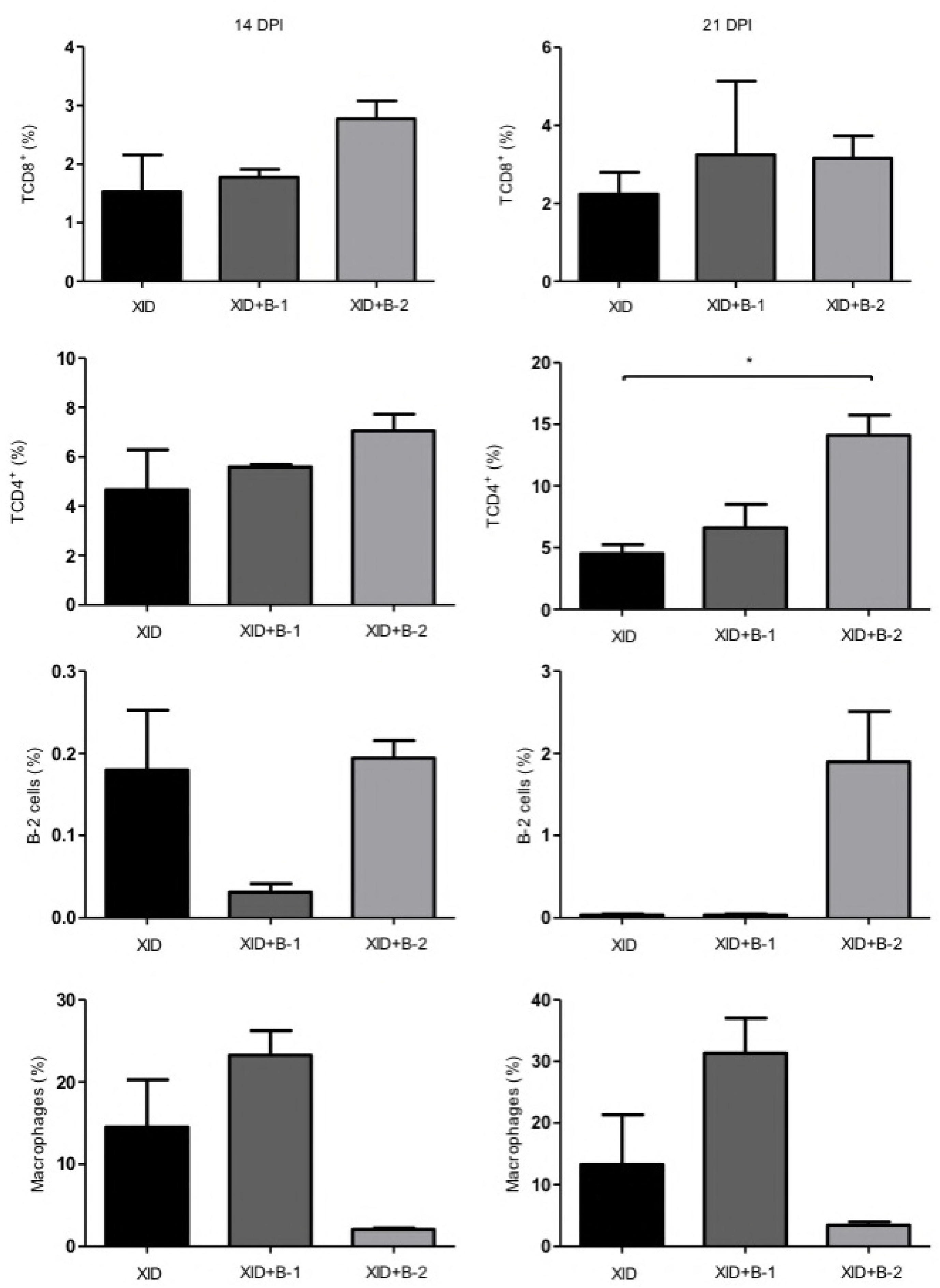
Evaluation of CD8^+^ and CD4^+^ T lymphocytes, B-1 and B-2 cells and macrophages in the spleen of XID, XID+B-1 and XID+B-2 mice immunosuppressed with cyclophosphamide and infected with *E. cuniculi* at 14 and 21 days post-infection (DPI). One-way variance analysis ANOVA with Bonferroni multiple comparisons post-test showed *p <0.5.

### *E. cuniculi* infection increased the levels of proinflammatory cytokines in XID mice

Cytokines dosage from the serum showed that *E. cuniculi* infection increased the levels of interferon-gama (IFN-γ), tumor necrosis factor (TNF-α) and interleukine – 6 (IL-6) in XID mice compared to its control and BALB/c groups (Fig. 12). Moreover, higher levels of these cytokines were observed in XID mice than those that received the adoptive transfer of B-1 and B-2 cells (Fig. 13). There was no statistical significance between IL-4 cytokine dosage from all groups (data not shown). IL-2, IL-10 and IL-17 cytokines were not detected.

**Figure 12.**
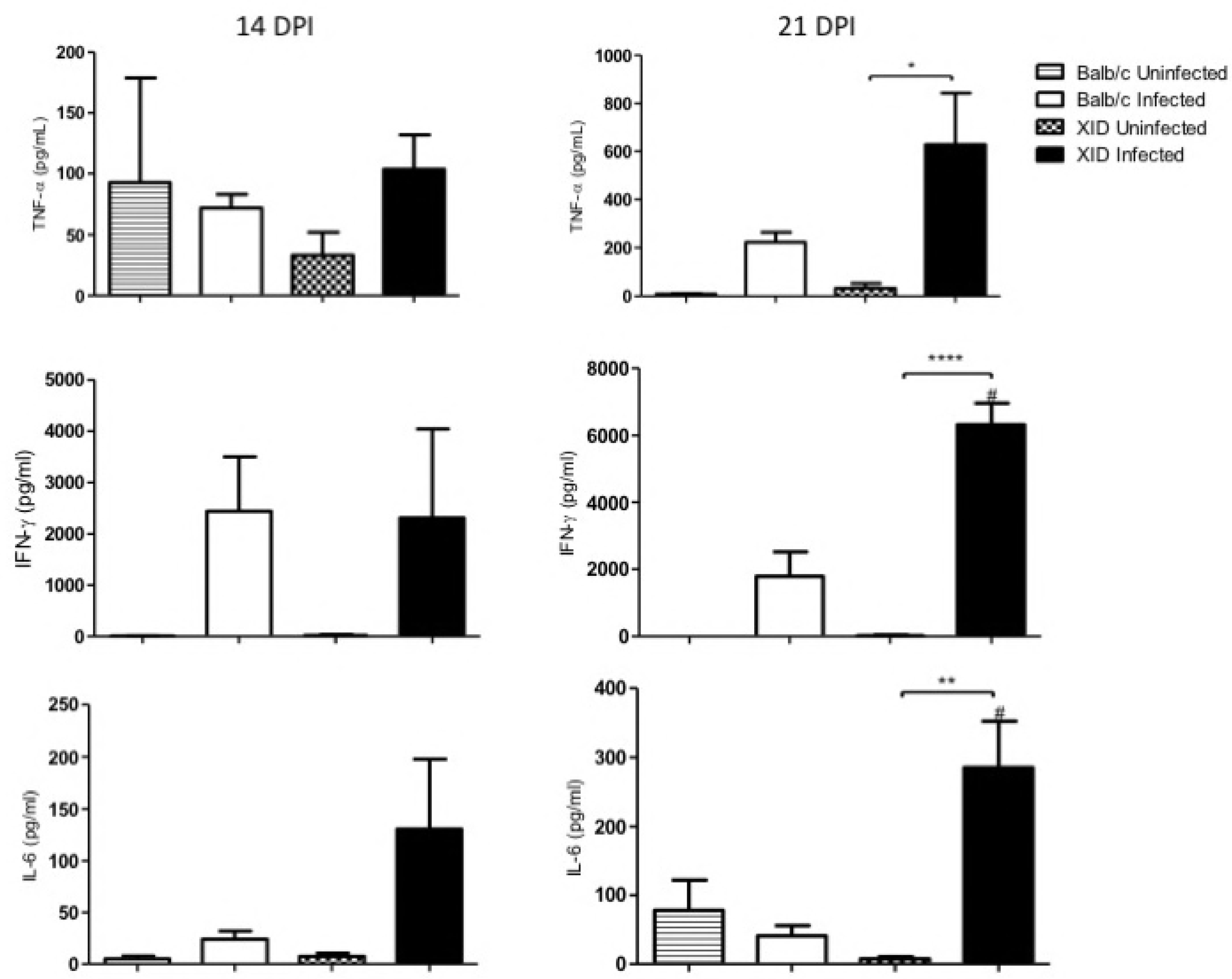
Evaluation of TNF-α, IFN-γ and IL-6 cytokines in the serum samples of BALB/c and XID mice immunosuppressed with cyclophosphamide and infected or not with *E. cuniculi* at 14 and 21 days post-infection (DPI). One-way variance analysis ANOVA with Bonferroni multiple comparisons post-test, *p<0.5; **p<0.01; ****p <0.001. Unpaired t-test, #p <0.01 in the comparison of BALB/c.

**Figure 13.**
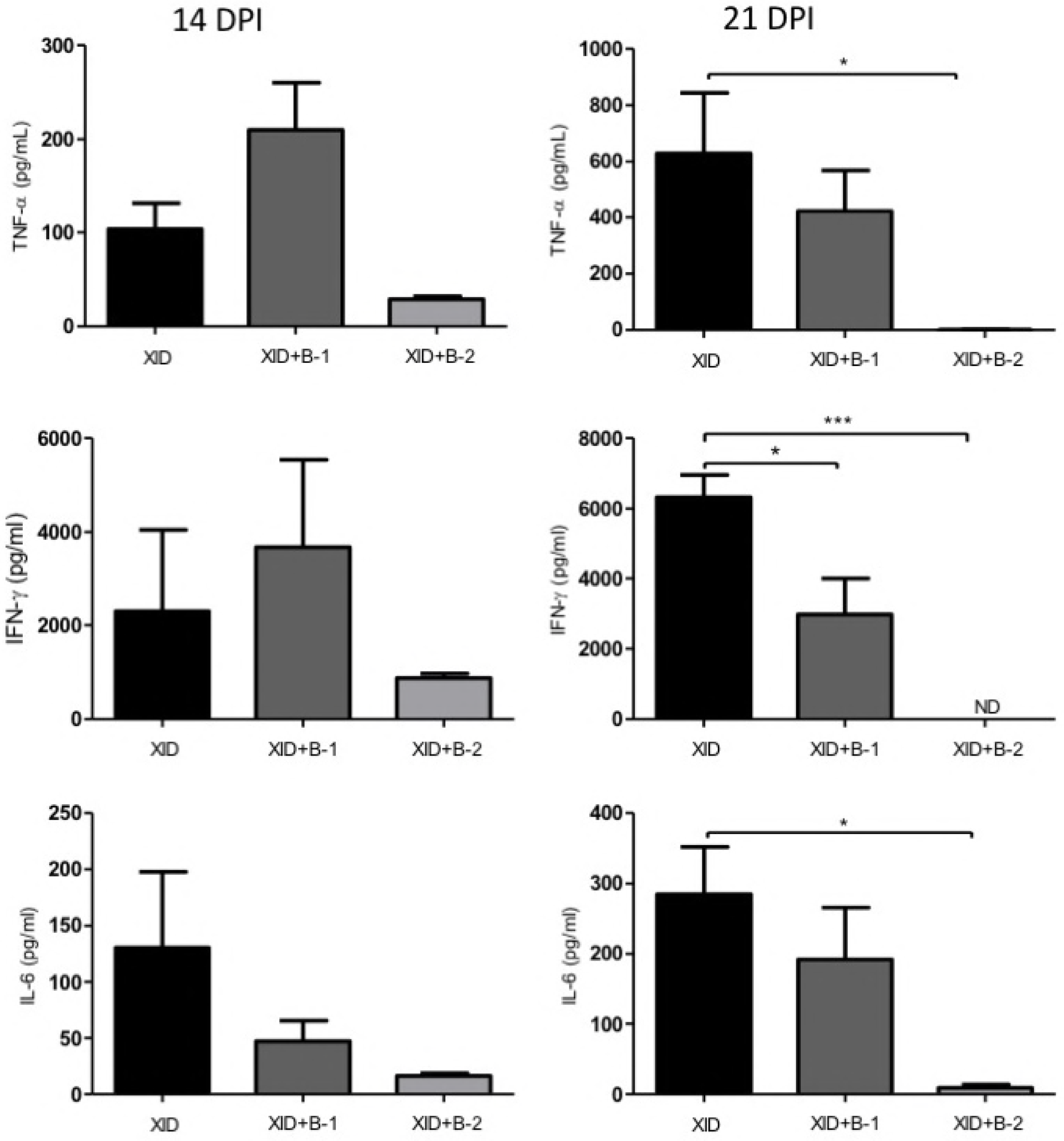
Evaluation of TNF-α, IFN-γ and IL-6 cytokines in the serum samples of XID, XID+B-1 and XID+B-2 treated with cyclophosphamide and infected with *E. cuniculi* at 14 and 21 days post-infection (DPI). One-way variance analysis ANOVA with Bonferroni multiple comparisons post-test revealed *p <0.5; ***p <0.001.

## Discussion

For the maintenance of the body integrity against a variety of pathogens, the immune system has different cell types. B cells are important in both innate and adaptive immune responses. Depending on the pathogen and infection route, subtypes of B cells, follicular or innate, differentiate specifically, which may be dependent or independent of T cells. Follicular B cells are specialized in the adaptive immune response, by mostly recognizing proteins, whereas B cells of the marginal zone (MZ) and B-1 cells support the innate immune response against non-protein antigens. Moreover, follicular B cells, localized in peripheric lymphoid organs, express specific receptors (BCR), which interact with antigens and became able to activate CD4 T lymphocytes, starting a T dependent immune response [26]. B cells of the marginal zone and B-1 cells express unspecific BCR and innate receptors which combined coordinate signals to B cells. Upon activation, these cells differentiate in extrafollicular plasmocytes [27].

In the present study, results showed that B-1 and B-2 cells deficiency in XID mice together with cyclophosphamide treatment caused a more severe and spread encephalitozoonosis, which was similar the described to SCID and nude mice [28]. B-1 cells have phagocytic and microbicide capabilities [29] and are considered very important in the defense against *Coxiella burnetti*, by its ability to produce antibodies, cytokines and promote phagocytosis [30]. The lack of B cells increased the susceptibility of CBA/XID to *Cryptococcus neoformans* disseminated fungal disease [31]. Our previous study showed that XID mice infected with *E. cuniculi* and not immunosuppressed were less resistant against development of lesions, with a decreased immune response compared to BALB/c mice by intraperitoneal and oral route infections [8,9]. *In vitro* studies of the immune response to microsporidia show that antibodies exert an opsonization effect and may block parasite entry into non-phagocytic cells [32], also reduced the growth of *E. cuniculi* in the presence of peritoneal macrophages and resulted in increased phagocytosis of spores [33,34]. The adoptive transfer of antibodies to nude or SCID mice was not sufficient to prevent the death of *E. cuniculi* infected animals [32], although, the effect of the antibodies on microsporidial infection in vivo was to prolong the survival of previously CD4^+^ reconstituted, perorally infected and intraperitoneally monoclonal antibody-treated SCID mice [34].

Although XID mice are frequently used to evaluate B-1 cells role they have a Bruton’s tyrosine kinase mutation, a protein responsible for the development of partial B cells population, therefore, they also have B-2 cells deficiency. In this study, we adoptively transferred WT B-2 cells to XID mice to evaluated B-2 cells role in the encephalitozoonosis. We observed that B-2 cells transfer decrease all immune populations in the peritoneal cavity and also downregulated all serum cytokines observed. In XID+B-1 mice, the opposite was observed, upregulation of peritoneal cells and serum cytokines. Into the spleen, B-2 populations increase in XID+B-2 and decrease in XID+B-1 and macrophages decrease in XID+ B-2 and increase in XID+B-1. In XID+B-2 mice which did not receive immunosuppression treatment, it was observed downregulation in peritoneal cells populations and higher resistance to disease development, with few clinical signs and histological lesions caused by *E. cuniculi* (article in preparation).

CD8^+^ T cells are critical for the survival of *E. cuniculi* infected host, since CD8^−/−^ mice die after inoculation of the pathogen, manifesting a marked and widespread disease characterized by lethargy, ascites, hepatitis, pneumonia and spleen lesions [6,13,15]. Cy-immunosuppressed XID mice showed as severe symptoms as CD8^−/−^ mice however peritoneal CD8^+^ and CD4^+^ T cells were increased in these animals, but not in BALB/c mice. We suggest a deficient response of T cells in XID mice. Previous studies showed B-1 cells ability in regulates other cells functions. B-1 cells may change the composition of the granuloma in the lungs caused by BCG in mice [35]. B-1 cells are also capable of increasing the migration of CD8+ T cells in grafts [36]. Moreover, peritoneal B-1 cells may also influence immune response by activating T cells without need migration to the targeted lymphoid organ [37]. Thus, we suggest that the absence of B-1 cells could affect the cytotoxic capacity of these lymphocytes in XID mice. Experiments are been performed in our lab to explain this hypothesis.

It should still be considered that the activity of peritoneal lymphocytes has not been fully understood. XID mice, although having a BALB/c background, may present a Th1 response similar to C57BL/6. In these mice, peritoneal macrophages activated by IFN-γ may promote suppression of peritoneal T cells by arginine consumption [38]. In contrast, BALB/c mice which present Th2 profile and low production of IFN-γ showed activation of peritoneal T cells in co-cultures with macrophages [38]. This phenomenon could clarify the response observed in XID mice in the present experiment, although the presence of the pathogen may promote the proliferation of CD8^+^ T lymphocytes, justifying the increase in their number as observed, the function of these cells could be suppressed by the presence of macrophages and high levels of IFN-γ, while in BALB/c the lower peritoneal population of CD8^+^ T cells associated with the milder infection could suggest an effective activation of these cells. Additionally, the increase in splenic CD4^+^ and CD8^+^ T cells of BALB/c mice suggests that these cells could be originated and activated in the spleen. A similar profile was identified in XID+B-1 mice, further suggesting a role of B-1 cells in the immune response mounting.

Macrophages increased in the peritoneum of infected XID mice. Pereira et al. (article in preparation) have demonstrated *in vitro* that the peritoneal macrophages from XID mice have predominantly M2 profile, high phagocytic and low microbicidal activities. Macrophages with M2 profile may favor the dissemination of the pathogen acting as a Trojan horse, explaining the severity of the disease in XID mice. In BALB/c mice, the dimensions of the areas of mononuclear inflammatory infiltrate into the liver secondary to infection were significantly larger than in XID, suggesting that this inflammatory response partially contained the development of the pathogen.

The microsporidia are among the most effective agents and developed for intracellular parasitism. After *E. cuniculi* infection, usually by the oral route, many mammals have a progressive disease after weeks or months. This infection is associated with high and persistent antibody in addition to a continuous inflammatory infiltration [39]. However, there are mice strains which are more susceptible to *E. cuniculi* infection, such as C57BL/6 [28] and as was observed in this and other studies, XID mice [8,9], but not the wild type BALB/c mice. It was demonstrated that C57BL/6 strain has a genetic alteration that compromises B lymphocyte function, so antibodies produced by these animals may have low efficiency [40]. In C57BL/6 mice, high levels of antibodies were identified after 14 days of experimental infection with *E. cuniculi*, but these animals showed significant lesions in the liver and brain associated with the presence of spores of the pathogen [41]. It is also emphasized that this mouse strain produces cytokine later than BALB/c mice, with distinct delay in the mobilization of the immune response [42].

the clinical symptoms were directly related to the severe inflammatory reaction observed in the cerebral tissue of infected rabbits

High levels of proinflammatory cytokines with Th1 profile were observed in the serum of XID than in BALB/c mice, which is in line with the clinical signs of more severe disease and corroborates the hypothesis of a profile similar to the C57BL/6 mice. Among these cytokines, IFN-γ, TNF-α, IL-6 were identified, which was the same cytokines observed previously from *E. cuniculi* infected mice but not Cy-immunosuppressed [43,44]. Th1 cytokines are important against Microsporidia, as previously shown in *in vivo* and *in vitro* model [43,44]. IFN-γ may also increase and plays important role in immunosuppressed mice [43]. In diabetic mice immunosuppressed with Cy IFN-γ was increased but not in diabetic non-immunosuppressed mice, suggesting an immunomodulatory effect of Cy in this cytokine production during encephalitozonosis [7]. The results herein presented showed that *E. cuniculi* infection together Cy immunosuppression increased IFN-γ both in XID and BALB/c mice. Cy is associated with increased serum IFN-γ and other proinflammatory cytokines [45].

We reinforce that infected XID mice had more severe encephalitozoonosis than BALB/c mice. Despite the immunosuppressive activity of Cy, the peritoneal populations of T-lymphocytes, macrophages and proinflammatory cytokines in infected XID mice increased, although splenic populations remained similar to uninfected controls. On the other hand, in BALB/c mice, we observed that splenic populations of CD4^+^ and CD8^+^ T lymphocytes increased, indicating the spleen as an important local for recruitment of the immune response to the Microsporidia. In conclusion, the Cy treatment allowed us to observe a more severe disease and evidenced differences between the immune response of BALB/c and XID mice. In addition, B-1 and B-2 cells transfer demonstrated the importance of both these populations in resistance to encephalitozoonosis. Differently, B-1 cells act modulating the immune system and although the role of B-2 cells is still unclear, we hypothesized that the production of antibodies by these cells could play an important role in the resolution of the disease and its should be deeply investigated.

## Materials and Methods

### Animals

Isogenic, female, with six to eight weeks old, *specific pathogen free*, BALB/c and BALB/c XID mice were obtained from the “Centro de Desenvolvimento de Modelos Experimentais para Biologia e Medicina” (CEDEME) from Federal University of Sao Paulo (UNIFESP, in Portuguese). All animals were divided in groups and kept in sterilized isolators at the animal facility at Paulista University (UNIP), under controlled temperature and humidity with water and food *ad libitum*.

### Ethics Statement

All experiments involving research animals were performed in accordance with the recommendations outlined in the Guide for the Care and Use of Laboratory Animals. All research animal protocols were approved by the institutional animal use (Comissão de Ética no Uso de Animais – CEUA) at Universidade Paulista - UNIP as part of the protocol numbers 138/12 and 385/15.

### B-2 cells adoptive transfer

XID mice were adoptively transferred with B-2 cells from the spleen of BALB/c mice [46]. After tissue dissociation, lysis of red blood cells and centrifugation, a pellet was incubated with anti-CD16/32 for blocking Fc receptors, for 15 minutes at 4°C. Then, cells were washed with phosphate buffered solution (PBS, Sigma, St Louis, MO, EUA) with1% Bovine Serum Albumin (BSA, Sigma, St Louis, MO, EUA) (PBS-BSA 1%). Next, cells were incubated with Phycoerythrin (PE) Cyanin 7 (Cy7)-conjugated rat anti-mouse CD19 and PE-conjugated rat anti-mouse CD23 antibodies for 20 minutes at 4°C for labelling of B-2 cells. Subsequently, cells were sorted in a cytometer *“cell sorter*” FACSAria II at UNIFESP. The population of lymphocytes was chosen based on *forward scatter* FSC versus *side scatter* SSC and the B-2 cells populations were selected according to the phenotype CD19^+^CD23^+^. Finally, centrifugated B-2 cells were resuspended in PBS and B-2 cells suspension (1×10^6^ cells in 200 μL) were injected intraperitoneally in XID mice seven days prior to day zero of infection, constituting XID+B-2 mice.

### B-1 cells adoptive transfer

XID mice were adoptively transferred with B-1 cells following adapted protocol from Almeida *et al*. (2001). Peritoneal cells from BALB/c mice were sampled by successive washes with 2 ml of RPMI-1640 (Sigma, St. Louis, MO. USA) and cultured for 40 minutes at 37°C with 5% CO_2_ atmosphere. Subsequently, the supernatant was discarded and cells that were adherent were washed and reincubated with RPMI plus10% bovine fetal serum (BFS) (R10) under the same conditions for further five days. Then, the supernatant was collected, centrifuged and resuspended in PBS. Finally, centrifugated B-1 cells enriched population were resuspended in PBS and were injected intraperitoneally (1×10^6^ cells in 200 μl) in XID mice seven days prior to day zero of infection, constituting XID+B-1 mice.

### Pathogen and experimental infection

*Encephalitozoon cuniculi*, spores, of genotype 1 (bought from Waterborne^®^ Inc. New Orleans, LA, USA) were grown in Rabbit Kidney (RK)-13 cells in the laboratory of Cell Culture at Paulista University. RK cells were kept in Eagle media supplemented with 10% BFS, Gibco, Grand Island, NY, EUA), plus 10% of non-essential amino acids, 10% pyruvate, and gentamicin (20 mg/ml) and incubated in 5% CO_2_ at 37°C. Supernatant was sampled every seven days, centrifugated for 20 minutes at 500*g* to obtain spores. *E. cuniculi* spores counting was done with a Neubauer’s chamber. All infected mice received intraperitoneal injection of 1×10^7^ *E. cuniculi* spores.

### Cyclophosphamide (Cy) treatment

All animals of infected and uninfected groups were treated with Cy (100 mg/kg, twice week). Treatment started at the day of infection until 14 and 21 days post infection (DPI).

### Necropsy and tissue sampling

Animals were euthanatized with of ketamine (100 mg/ml), xylazine (20 mg/ml) and fentanyl (0.05 mg/ml) at 14 and 21 DPI. Firstly, the blood was sampled from the heart puncture for serum sampling and cytokines dosage; secondly, a peritoneal wash was performed and the spleen was sampled and processed for flow cytometry (see below); and thirdly tissue samples (liver, spleen, lungs, intestine) were sampled for histopathology and TEM.

### Histological and morphometric analysis

Samples of the liver, lungs, small intestine and spleen were fixed in 10% buffered formalin solution (pH 7.2–7.4). Tissue samples were routinely processed and embedded in paraffin to obtain tissue 4μm samples for hematoxylin and eosin staining. Inflammatory infiltrates in the liver were analyzed by *MetaMorph Software* (Molecular Devices, California, EUA). Ten random photomicrographs were taken and measured by pixels area, and the means of each group were used for statistical analysis.

### Fungal burden

We count *E. cuniculi* free spores in the peritoneal wash from each infected animal. The mean of each animal was calculated in spores/mL and statistically analyzed.

### Ultrastructural analysis by transmission electronic microscopy (TEM)

Samples of 1mm from the liver and spleen were from XID infected mice were fixed in glutaraldehyde at 2% in cacodylate buffer 0.2 M (pH 7.2) at 4°C for 10 hours, post fixed in buffered OsO4 at 1% for 2 hours, routinely processed, and embedded in Epon’s resin. Semithin cuts were contrasted with Toluidine blue and ultrathin cuts were doubled colored with aqueous uranyl and lead citrate. Finally, samples were observed in a TEM LEO EM 906 at 80 kV at Butantan Institute.

### Immunohistochemistry

Four μm sections were cut and mounted on slides with poly-L-lysine for better adhesion. The slides were then dewaxed in xylene for 40 min at 75 C and then rehydrated in graded alcohol and distilled water. Antigen retrieval was achieved by heating the slides at 90_C in a buffer solution of 0,01M sodium citrate, at pH 6.5, for 25 min in a steamer. Blockage of endogenous peroxidase activity was performed by incubation with hydrogen peroxide 1,75% in methanol for 5 min. and non-specific interactions were blocked with 5% skimmed milk for 5 min. The slides were incubated with a primary mouse monoclonal anti-pancytokeratin antibody (clone AE1/AE3, Biocare Medical, 1:400) for 30 min.; Mach 1 probe, for 30 min., Mach 1 polymer (both from Biocare Medical) for 30 min. Labelling was ‘visualized’ with 3,30-diaminobenzidine (Biocare Medical) and sections were counterstained with Harris’ haematoxylin (Hsu *et al* 1981). The healthy liver tissue was used as positive control.

### Phenotyping of immune cells from the peritoneal cavity and spleen

Peritoneal cells were obtained as described. The spleen was mechanically dissociated and filtered with a 70 μm cell strainer; red blood cells were removed by hemolytic buffer. After centrifugation, cells were washed with PBS and resuspended in 100 μl of PBS – BSA 1%. Each sample was then incubated at 4°C for 20 min with anti-CD16/CD32 antibody. After incubation, cells were divided in two aliquots and ressupended in PBS-BSA 1%; and subsequently incubated with the monoclonal antibodies: *Allophycocyanin* (APC)- conjugated rat anti-mouse CD19, *Fluorescein Isothiocyanate* (FITC) or PE-conjugated rat anti-mouse CD23, *Peridinin Chlorophyll Protein Complex* (PerCP)-conjugated rat anti-mouse CD4, FITC-conjugated rat anti-mouse CD8, PE-conjugated rat anti-mouse F4/80 and *Pacific Blue* or APC-Cy 7-conjugated rat anti-mouse CD11b (BD-Pharmingen, San Diego, CA). Finally, cell suspensions were analyzed in the flow cytometer FACS Canto II (BD Biosciences, Mountain View, CA, USA). Cells were characterized according to their phenotypes, as shown in Table 1, and analyzed with the software FlowJo (FlowJo LLC, Data Analysis Software, Ashland, OR).

**Table 1.** Panel of cell surface markers used in this study for immunophenotyping.

| Phenotype | Markers |
| --- | --- |
| B-1cells | CD23<sup>-</sup> CD19<sup>+</sup> |
| B-2 cells | CD23<sup>+</sup> CD19<sup>+</sup> |
| TCD4<sup>+</sup> cells | CD19<sup>-</sup> CD4<sup>+</sup> |
| TCD8<sup>+</sup> cells | CD19<sup>-</sup> CD8<sup>+</sup> |
| Macrophages | CD19<sup>-</sup> CD11b<sup>+</sup> F4/80<sup>+</sup> |

### Cytokines quantification

The serum was obtained from the blood and kept at −80°C until thawed for cytokines dosage by flow cytometry, using CBA Mouse Th1/Th2/Th17 Cytokine Kit (BD Biosciences, CA, EUA), according to the manufacturer’s instructions. 25 μl of each sample, 25 μl of mixed beads containing specific capture antibodies and 25 μl of PE-conjugated detection antibody were incubated for 2 hours at room temperature in the dark. Subsequently, samples were centrifugated, washed and resuspended for analysis by two colors flow cytometry by using FACS Canto II (BD Biosciences, Mountain View, CA, USA). All analysis was done by using the software FCAP Array 3.0.

### Statistical analysis

Comparisons were made by variance analysis (ANOVA) of one or two ways and Tukey’s or Bonferroni multiple comparisons posttests. All values were showed as mean ± standard error mean (SEM) with significance of α=0,05 (p<0,05). All graphics were made at *“GraphPad Prism*” version 5.0 for Windows ^®^ (GraphPad Software Inc, La Jolla, CA, EUA).

## Acknowledgments

We thank Magna Maltauro Soares (Instituto Butantan) for histopathological processing.

